# Soluble αβ-tubulins reversibly sequester TTC5 to regulate tubulin mRNA decay

**DOI:** 10.1101/2024.04.12.589224

**Authors:** Alina Batiuk, Markus Höpfler, Ana C. Almeida, Deryn Teoh En-Jie, Oscar Vadas, Evangelia Vartholomaiou, Ramanujan S. Hegde, Zhewang Lin, Ivana Gasic

## Abstract

Microtubules, built from heterodimers of α- and β-tubulins, control cell shape, mediate intracellular transport and power cell division. The concentration of αβ-tubulins is tightly controlled through a post-transcriptional mechanism involving selective and regulated degradation of tubulin-encoding mRNAs. Degradation is initiated by TTC5, which recognizes tubulin-synthesizing ribosomes and recruits downstream effectors to trigger mRNA deadenylation. Here, we have investigated how cells regulate TTC5 activity. Biochemical and structural proteomic approaches reveal that under normal conditions, soluble αβ-tubulins bind to and sequester TTC5, preventing it from engaging nascent tubulins at translating ribosomes. We identify the flexible C-terminal tail of TTC5 as a molecular switch, toggling between soluble αβ-tubulin-bound and nascent tubulin-bound states. Loss of sequestration by soluble αβ-tubulins constitutively activates TTC5, leading to diminished tubulin mRNA levels and compromised microtubule-dependent chromosome segregation during cell division. Our findings provide a paradigm for how cells regulate the activity of a specificity factor to adapt posttranscriptional regulation of gene expression to cellular needs.

## Introduction

Built from heterodimers comprising α- and β-tubulin proteins (αβ-tubulins hereafter), microtubules are core eukaryotic cytoskeletal elements^1^. Cells rely on microtubules for the organization of their internal contents, motility, and division^2^. To execute their roles effectively, cells must tightly regulate the spatial distribution and dynamic properties of microtubules^3^, achieved through numerous regulatory pathways. Foremost among these is microtubule dynamic instability manifested in consecutive phases of growth through the addition, and shrinkage through the loss of αβ-tubulins^4^. By regulating dynamic instability through microtubule and tubulin-binding proteins^5,6^, cells can use microtubule growth and shrinkage to power various physical processes such as chromosome segregation during cell division.

Microtubule dynamics are critically dependent on the concentration of soluble αβ-tubulins. As their core building blocks, soluble αβ-tubulins directly impact microtubule nucleation, polymerization, and dynamic properties^7,8^. Cells therefore tightly regulate the availability of soluble αβ-tubulins through a feedback loop that restricts the biosynthesis of new tubulin when αβ-tubulins are in surplus^9,10^. Termed tubulin autoregulation, this pathway involves the selective degradation of tubulin-encoding mRNAs in a translation-dependent reaction^9,11–13^. The mechanism involves a ribosome-binding factor termed TTC5 (tetratricopeptide protein 5), which selectively recognizes the N-terminal sequences of nascent α- and β-tubulins at the ribosomal exit tunnel^14^. TTC5 recruits the adaptor protein SCAPER (S-Phase Cyclin A Associated Protein in the ER), which recruits the large deadenylase complex CCR4-NOT (Carbon Catabolite Repression—Negative On TATA-less), initiating tubulin mRNA deadenylation and decay^15^.

Mutations in TTC5 or SCAPER associated with complete or near-complete loss of protein have been linked to tubulinopathies—a class of neurodevelopmental disorders arising from mutations in tubulins^16–21^. Disruption of tubulin autoregulation in cultured cells compromises mitotic fidelity^14,15^, a phenotype frequently attributed to aberrant microtubule dynamics^22,23^. This phenotype is seen with mutations that perturb TTC5 recognition of the ribosome, recognition of nascent tubulins, recruitment of SCAPER, or SCAPER-mediated recruitment of CCR4-NOT^14,15^. Thus, the ribosome-proximal molecular events in tubulin autoregulation that culminate in mRNA decay are now generally well established^14,15^. By contrast, the mechanisms that control deployment of this mRNA decay machinery remain unknown. In this study, we focus on how cells control the activity of TTC5, which serves as both the specificity factor and the most upstream component of ribosome-associated mRNA decay^14,15^.

## Results

### Soluble αβ-tubulins can directly inhibit TTC5 activity

TTC5 binding to tubulin-synthesizing ribosomes initiates tubulin mRNA decay^14^. Because tubulin mRNA degradation is thought to be modulated by the cell in response to perceived tubulin need, we speculated that TTC5 abundance, TTC5 localization, or TTC5 engagement of ribosome-nascent chain complexes (RNCs) is likely to be regulated (**Figure 1A**). Immunoblotting showed that TTC5 protein levels remain constant before and after acute microtubule destabilization by colchicine (COL), combretastatin A4 (CA4), or nocodazole (NOC, **Figure S1A**), each of which triggers tubulin mRNA degradation^9,24^. Furthermore, a model where microtubule-sequestered TTC5 is released upon depolymerization seems unlikely because GFP-TTC5 (fully competent for tubulin autoregulation^15^) is diffusely cytosolic without an obvious microtubule localized population (**Figure S1B**). Finally, simply overexpressing TTC5 in cells is insufficient to trigger tubulin mRNA degradation (**Figure S1C**) unless the autoregulation pathway is triggered by microtubule destabilization^14^. These results suggest that TTC5 is constitutively present in the cytosol, but does not initiate tubulin mRNA degradation under normal steady-state conditions.

**Figure 1.**
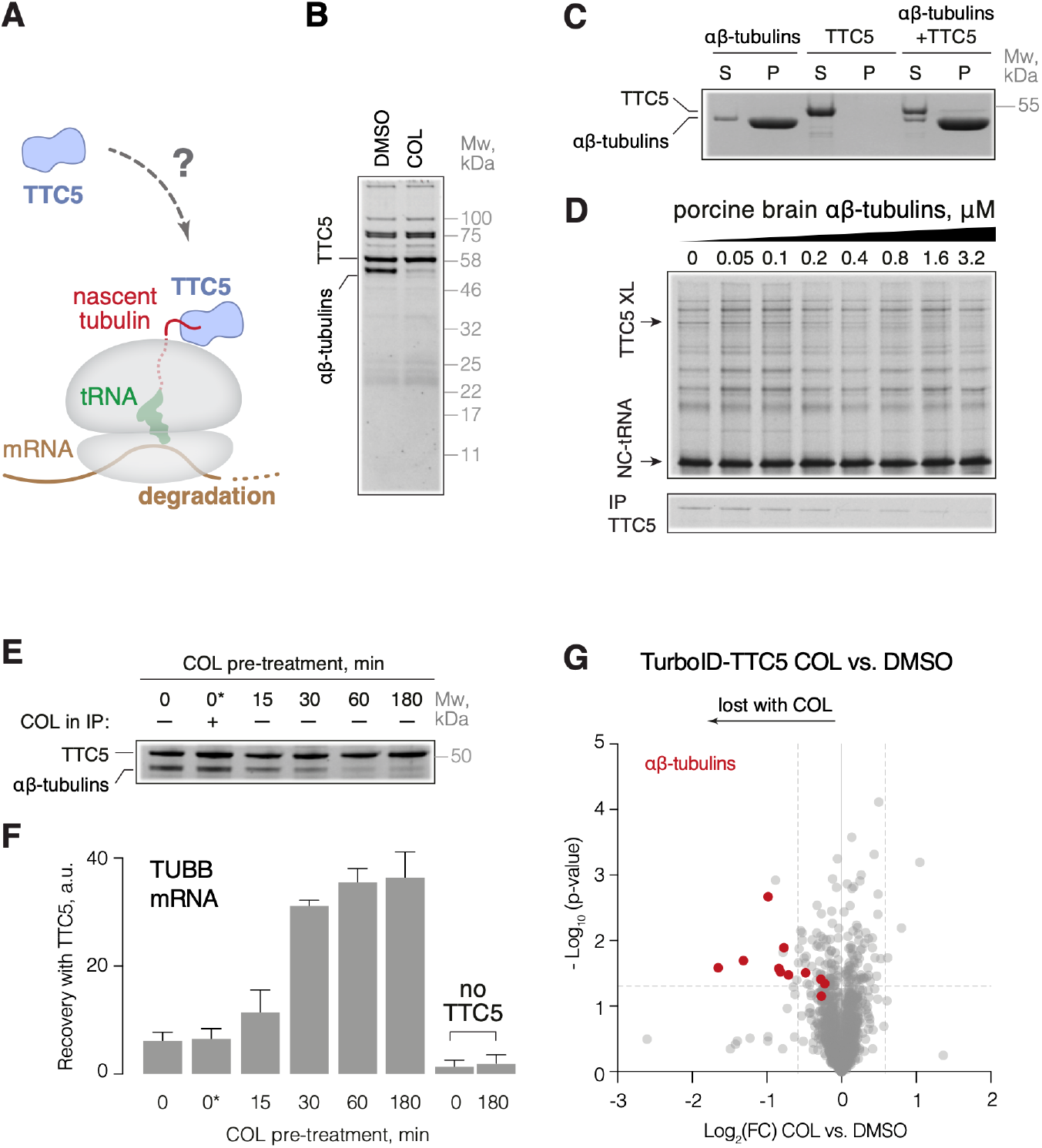
Soluble αβ-tubulins reversibly repress TTC5 to regulate its activity. **(A)** Schematic representation of the first step of the tubulin autoregulation pathway. **(B)** Recombinant Strep-tagged TTC5 was mixed with lysate from untreated or colchicine-treated (COL, for 3 hours) TTC5 knockout HEK293 cells and recovered by affinity purification via the Strep tag. The sole differential interaction partner was a mixture of αβ-tubulins. **(C)** Binding of TTC5 to microtubules was analyzed by microtubule co-pelleting assay. TTC5 mostly remains in the soluble fraction (S), while microtubules are mostly in the pellet (P). **(D)** Nascent β-tubulin crosslinking assay in the presence of the indicated concentrations of soluble porcine brain αβ-tubulins. The TTC5 crosslink is indicated and verified by immunoprecipitation in the bottom panel. **(E)** TTC5 knockout HEK293 cells were pre-treated for the indicated times with colchicine and used to prepare lysates. One aliquot of each lysate was used for binding analysis to recombinant Strep-TTC5 as in panel B. One of the control samples included colchicine added after cell lysis (indicated as 0*). **(F)** The products coimmunoprecipitated with recombinant TTC5 in panel E were analyzed for α-(**Figure S1F**) and β-tubulin mRNAs by quantitative RT-PCR (mean ± SD from three independent replicates). **(G)** Proximity labeling using TurboID fused to TTC5 and expressed in TTC5 knockout HEK293 cells, followed by enrichment of biotinylated proteins and quantitative mass spectrometry. Data from cells treated with colchicine were normalized to DMSO control and plotted as Log2 fold-change (Log2(FC)). Dashed lines represent 1.5-fold change and 0.05 p-value. Highlighted in red are tubulins.

Previously, cytosol extracted from untreated cells, but not colchicine-treated cells, was proposed to contain an inhibitor of TTC5 activity^14^. In this experiment, an *in vitro* assembled complex between TTC5 and tubulin-synthesizing RNCs could be disrupted by cytosol from cells growing under normal conditions but not from cells undergoing active tubulin mRNA decay induced by colchicine treatment^14^. Speculating that this putative inhibitory factor might act directly on TTC5, we used recombinant immobilized TTC5 to identify interaction partners from the cytosol of untreated versus colchicine-treated TTC5 knockout cells. The only differentially interacting protein recovered stoichiometrically with TTC5 from untreated cell lysates was identified by mass spectrometry as a mixture of α- and β-tubulins (**Figure 1B**).

Because the cell lysates were prepared on ice, both the untreated and colchicine-treated cytosol had equal amounts of depolymerized αβ-tubulins, with little or no intact microtubules (whose polymerization is temperature-dependent). Indeed, no appreciable TTC5 co-sedimented with polymerized purified porcine brain αβ-tubulins (**Figure 1C**), consistent with an absence of GFP-TTC5 co-localization with microtubules in cells (**Figure S1B**). By contrast, soluble αβ-tubulins interacted with recombinant TTC5 with Kd of ∼0.5 μM (**Figure S1D**) and progressively inhibited TTC5-RNC interaction with an estimated Ki of ∼0.5 μM (**Figure 1D**). Thus, soluble αβ-tubulins at physiological concentrations (1-3 μM) are direct competitors of RNCs for binding to TTC5.

### TTC5 inhibition by αβ-tubulins is attenuated during autoregulation

The capacity of soluble αβ-tubulins to bind immobilized TTC5 was progressively lost over the course of 60 minutes when cells were pre-treated with colchicine prior to preparation of cytosol (**Figure 1E**). Importantly, post-lysis addition of colchicine to cytosol prepared from untreated cells had no effect on the αβ-tubulin-TTC5 interaction. Similarly, purified porcine αβ-tubulins showed no difference in their TTC5 interaction regardless of any pre-incubation with colchicine (**Figure S1E**).

Thus, colchicine treatment of live cells, but not cytosol or purified tubulins, triggers a yet-unidentified change to the capacity of soluble αβ-tubulins to interact with TTC5. This loss of interaction with αβ-tubulins correlates with the disappearance of a TTC5-inhibitory activity in cytosol as monitored using *in vitro* assays, and further correlates with the initiation of tubulin mRNA decay in cells^9^.

Colchicine-triggered loss of a TTC5 inhibitor was further supported by the finding that recombinant TTC5 could selectively pulldown tubulin-encoding mRNAs from the cytosol of colchicine-pretreated TTC5 knockout cells (**Figures 1F, S1F**). Roughly six- to eight-fold less tubulin mRNA was pulled down by TTC5 from the cytosol of untreated cells or cytosol treated with colchicine after cell lysis. Notably, the 60-minute time course of colchicine-triggered tubulin mRNA recovery by TTC5 mirrored the loss of interaction between αβ-tubulin and TTC5. Total tubulin mRNA levels remained unchanged throughout these treatments as expected for TTC5 knockout cells (**Figure S1G**). A similar colchicine-triggered shift of TTC5 interaction partners was seen in proteomic analysis of GFP-TTC5 pulldowns before and after colchicine treatment. Here, we observed a markedly reduced recovery of several tubulin isotypes and an increased recovery of ribosomal proteins and SCAPER upon treatment with colchicine (**Figure S1H**). Similar losses of TTC5-tubulin interactions were seen with nocodazole-mediated microtubule destabilization (**Figure S1I**).

To directly monitor the autoregulation-triggered disruption of the tubulin-TTC5 interaction in living cells, we carried out proximity labeling using the promiscuous biotin ligase TurboID fused to TTC5. Quantitative mass spectrometry of biotinylated proteins revealed that tubulins were strongly proximal to TTC5 in untreated cells but substantially less in cells treated with colchicine (**Figures 1G, S1J, S1K**). Thus, TTC5 is engaged with soluble αβ-tubulins in cells and in vitro. The clear anti-correlation of the αβ-tubulin-TTC5 interaction versus the capacity of TTC5 to engage tubulin-synthesizing RNCs and initiate mRNA decay argues that soluble αβ-tubulins are potent repressors of TTC5 activity. This repressive interaction is progressively lost upon conditions that trigger tubulin mRNA decay, indicating that a key control point in tubulin autoregulation is reversible sequestration of TTC5 by soluble αβ-tubulins.

### C-terminal tail of TTC5 acts as a molecular switch

To identify TTC5 domains required for repression by αβ-tubulins, we used hydrogen-deuterium exchange-based structural mass spectrometry (HDX-MS). In HDX-MS experiments, amide-bond hydrogens in dynamic regions of proteins are exchanged for deuterium, which can be monitored by mass spectrometry^25^. TTC5 in isolation showed high deuteration in loops connecting alpha-helices of its tetratricopeptide repeats, and a near complete deuteration in its ∼20 amino acid C-terminal tail (**Figures 2A, 2B, S2A**) suggesting that this region is devoid of secondary structure elements. Of these deuterated sites, only the C-terminal tail (residues ∼420-440) showed marked protection from deuteration in the presence of αβ-tubulins. Modest but specific reductions in deuteration were also seen at residues 127-164, 193-209, and 367-379, regions that are all on the same face of TTC5 (**Figures 2C, 2D**, **S2B**). These observations suggest that TTC5 interaction with αβ-tubulins buries or otherwise alters these regions of TTC5, particularly its C-terminal tail.

**Figure 2.**
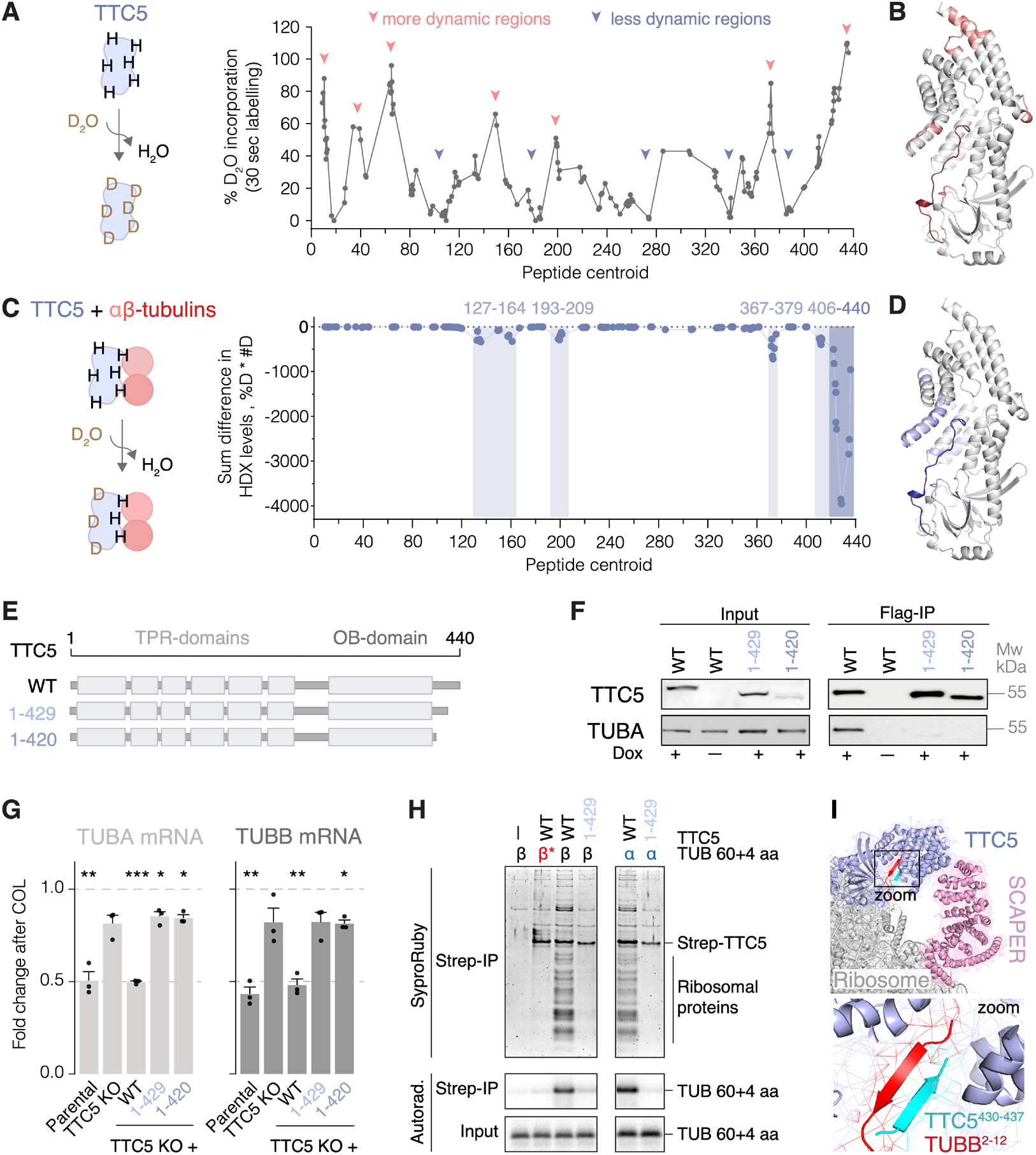
: C-terminal tail mediates interaction with mature and nascent tubulin. **(A)** Schematic of HDX-MS approach (left). Deuteration profile of human recombinant Strep-TTC5 over the full protein length identifies C-terminal domain as highly flexible or dynamic (right). Data are represented as means over three independent replicates. **(B)** Deuteration profile mapped onto the AlphaFold2-predicted structure of human TTC5 (AF-Q8N0Z6-F1)^26^. **(C)** Schematic representation of HDX-MS approach on the αβ-tubulin/TTC5 complex (left), and differential deuteration of human recombinant Strep-TTC5 upon incubation with porcine brain tubulin in deuterated water for 5 minutes (right). Data are presented as a sum of difference in hydrogen/deuterium exchange (% deuterons * number of deuterons) in the TTC5 + αβ-tubulins versus TTC5 alone samples in three independent replicates. Shaded areas highlight regions on TTC5 that show significantly lower deuteration upon binding to αβ-tubulin compared to unbound state. **(D)** Differential deuteration profile mapped onto the AlphaFold2-predicted structure of human TTC5. **(E)** Schematic representation of the wild type and generated TTC5 mutants. **(F)** Indicated Flag-tagged TTC5 constructs were expressed in HeLa TTC5 knockout cells under a doxycycline-inducible promoter and affinity purified via the Flag-tag. Coimmunoprecipitated interactors were separated using SDS-PAGE and tubulins visualized using western blot. **(G)** Autoregulation assay with HeLa parental, TTC5 knockout, and the indicated Flag-TTC5 rescue cell lines. Data show the mean ± SD mRNA levels after colchicine treatment from three independent experiments. *, **, and *** indicate p < 0.05, p < 0.01, and p < 0.001, respectively, in unpaired, two-tailed Student’s t tests for each of the indicated cell lines with the DMSO-treated sample as reference. **(H)** 64-residue α- and β-tubulin (TUB 60+4 aa) nascent chains were produced in rabbit reticulocyte lysates in the presence of recombinant wild type (WT) or mutated Strep-TTC5. Strep-TTC5 and its associated proteins were enriched via the Strep tag. Interacting partners were separated by SDS-PAGE and visualized by SYPRO Ruby staining for total protein and autoradiography for the α- and β-tubulin nascent chains (TUB 60+4 aa). β* indicates a β-tubulin construct in which its TTC5-interacting MREI motif has been mutated to autoregulation-incompatible MHQV. **(I)** Close-up view of the AlphaFold2-predicted C-terminal domain of TTC5 (cyan) and nascent β-tubulin (red) forming a beta sheet, fitted into the experimental cryo-EM density from a recent study (PDB: 8BPO)^15^.

The TTC5 C-terminal tail is required for its interaction with αβ-tubulins as seen in pulldown experiments with two truncated TTC5 mutants, TTC5^1-429^ and TTC5^1-420^ (**Figure 2E, 2F**). Similar results were obtained in cells using proximity labeling. Immunostaining for biotinylated proteins revealed that tubulins were substantially less biotinylated in TTC5 KO cells re-expressing Turbo-ID fused to either TTC5^1-429^ or TTC5^1-420^ compared to wild type TTC5 (**Figure S2C**). In the absence of sequestration by αβ-tubulins, TTC5^1-429^ or TTC5^1-420^ were expected to constitutively engage tubulin RNCs and trigger tubulin mRNA decay. Unexpectedly, TTC5^1-429^ and TTC5^1-420^ re-expressed in TTC5 knockout cells failed to degrade tubulin mRNA even upon colchicine stimulation (**Figure 2G, S2D**), suggesting that TTC5’s C-terminal tail has a functional role in mRNA decay.

To identify the step where the C-terminal tail plays a role, we analyzed the TTC5 interaction with *in vitro* produced RNCs of α- or β-tubulins. TTC5^1-429^ was unable to effectively recover tubulin RNCs in a pulldown assay (**Figure 2H**), indicating that the C-terminal tail is critical for recognition of tubulin-synthesizing RNCs. This was surprising because this tail was not modeled in the initial cryo-electron microscopy (EM) structure of tubulin RNCs engaged by TTC5^14^. Remarkably, an AlphaFold2^26^ prediction with full length TTC5 and the N-terminal region of β-tubulin showed the C-terminal tail of TTC5 (residues 431-436) forming an anti-parallel beta sheet with residues 4-8 in nascent β-tubulin, both of which are housed in TTC5’s substrate-binding groove (**Figures S2E, S2F**). We then re-inspected the improved cryo-EM map from a more recent study^15^ and found that the density in the binding groove is consistent with a two-stranded beta sheet (**Figures 2I, S2E, S2G**). The snug fit of the N- and C-terminal tails of nascent tubulin and TTC5, respectively, into TTC5’s groove appears to be important for stable association of TTC5 with tubulin-translating ribosomes.

Taken together, these results identify the flexible and unstructured C-terminal tail as having a critical role in two independent steps of the tubulin autoregulation pathway. Under steady-state conditions, this tail engages with and stabilizes an interaction between TTC5 and αβ-tubulins in a repressive complex. Although the molecular details of this interaction will require structural analysis, a direct role for the C-terminal tail is supported by both HDX-MS and the consequences of its deletion *in vitro* and in cells. Under autoregulation conditions, the same C-terminal tail forms a complex with nascent tubulin inside the binding groove of TTC5 at the ribosome. Thus, TTC5’s C-terminal tail acts as a molecular switch, toggling between a repressive complex with soluble αβ-tubulins and an activated complex with nascent tubulin on the ribosome.

### Loss of binding to αβ-tubulins constitutively activates TTC5

The N-terminal segment of nascent α-tubulin or β-tubulin engaged by TTC5 on the ribosome is buried in αβ-tubulins^27^. This suggests that although the C-terminal tail of TTC5 is involved in both interactions, the molecular details are likely to differ. Furthermore, the TTC5 interaction with αβ-tubulins probably involves other regions of TTC5 as indicated by the HDX-MS results. We therefore reasoned that it might be possible to identify TTC5 mutants that lack its repressive interaction with αβ-tubulin while preserving its activity on tubulin-translating ribosomes. Such a mutant would allow us to test the biological importance of the repressive interaction with αβ-tubulin directly and specifically.

Focusing first on the C-terminal tail we found that mutating V430 and T432 to glutamic acid (TTC5^VTEE^) resulted in near complete loss of binding to αβ-tubulins (**Figures 3A, 3B**). T432E alone was sufficient to mostly recapitulate this binding defect with αβ-tubulins as seen in pulldowns from cells (**Figures 3A, 3B**). In a second approach, we mutated various conserved surface residues in TTC5 (**Figure S3A**) and found by pulldown that TTC5^D175A^ was markedly reduced in its ability to bind αβ-tubulins (**Figures 3B, S3B**).

**Figure 3:**
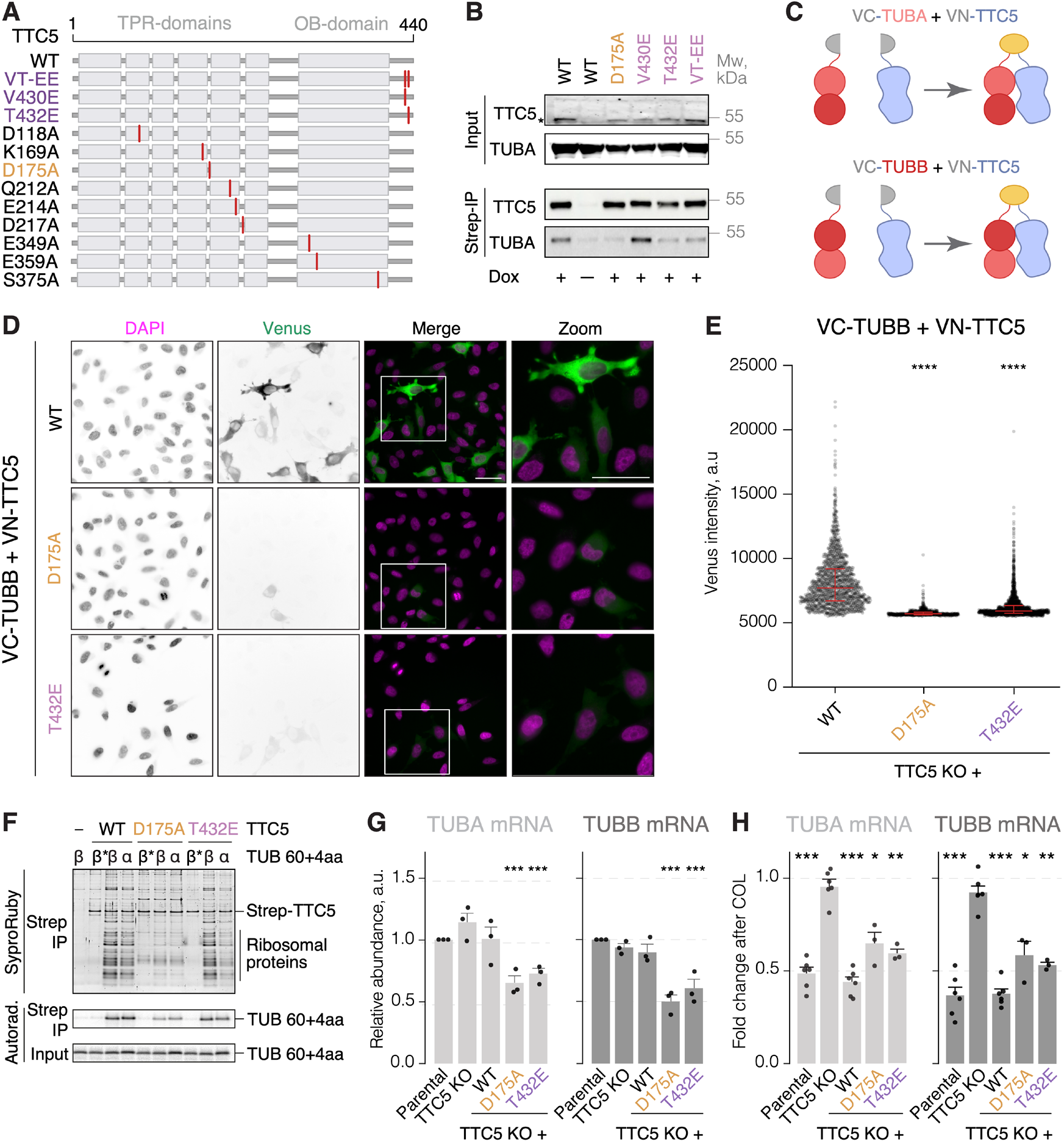
Loss of binding to αβ-tubulins constitutively activates TTC5. **(A)** Schematic representation of the wild-type (WT) and generated TTC5 mutants. **(B)** Indicated Strep-TTC5 constructs were expressed in HeLa TTC5 knockout cells and pulled down via Strep-tag. Bound αβ-tubulins were separated using SDS-PAGE and visualized by western blot. Asterisk indicates Strep-TTC5 band. **(C)** Schematic representation of the BiFC approach. **(D)** Representative live-cell images of the indicated VN-TTC5 and VC-TUBB constructs expressed in HeLa TTC5 knockout cells. Scale bar = 20 μm. **(E)** Fluorescence intensity of Venus in cell lines expressing the indicated BiFC constructs. Median and interquartile ranges of fluorescence intensities are depicted with red lines. Quadruple asterisks indicate p < 0.0001 in Mann-Whitney test for each of the indicated BiFC constructs with the BiFC constructs based on TTC5^WT^ as reference. **(F)** 64-residue α- and β-tubulin (TUB 60+4 aa) nascent chains were produced in rabbit reticulocyte lysates in the presence of recombinant WT or mutated Strep-TTC5. Strep-TTC5 and its associated proteins were subsequently enriched via the Strep tag. Interacting partners were separated by SDS-PAGE and visualized by SYPRO Ruby staining for total protein and autoradiography for the α- and β-tubulin nascent chains (TUB 60+4 aa). β* indicates a β-tubulin construct in which its TTC5-interacting MREI motif has been mutated to autoregulation-incompatible MHQV. **(G)** Relative α- and β-tubulin mRNA levels in HeLa parental, TTC5 knockout, and the indicated Strep-TTC5 rescue cell lines, normalized to a housekeeping transcript and the parental cell line. Data show the mean ± SD from three independent experiments. Single, double, and triple asterisks indicate p < 0.05, p < 0.01, and p < 0.001, respectively, in unpaired, two-tailed Student’s t tests for each of the indicated cell lines with the parental cell line as reference. **(H)** Autoregulation assay with HeLa parental, TTC5 knockout, and the indicated Strep-TTC5 rescue cell lines. Data show the mean ± SD mRNA levels after colchicine treatment, from at least three independent experiments. Single, double, and triple asterisks indicate p < 0.05, p < 0.01, and p < 0.001, respectively, in unpaired, two-tailed Student’s t-tests for each of the indicated cell lines with the DMSO-treated sample as reference.

TTC5^D175A^ and TTC5^T432E^ were analyzed for their ability to interact with αβ-tubulins in live cells using bimolecular fluorescence complementation (BiFC)^28–30^. In these experiments, the N-terminal fragment of the yellow fluorescent protein (Venus^1-172^, VN) was appended to TTC5, and the C-terminal fragment of the yellow fluorescent protein (Venus^156-238^, VC) was fused to α-(TUBA1B) or β-tubulin (TUBB, **Figure 3C, S3C**). The two fragment-fused proteins were expressed at equal levels in TTC5 KO cells from a single tandem open reading frame separated with a self-cleaving peptide (P2A). Both TTC5^D175A^ and TTC5^T432E^ BiFC constructs showed reduced fluorescence intensity relative to wild type TTC5 when paired with either α- or β-tubulin (**Figures 3D, 3E**, **S3D, S3E**). The expression levels of TTC5 mutants were comparable to wild type TTC5, indicating that the reduced fluorescence signal was due to reduced interaction (**Figure S3F**). Similar conclusions were reached from proximity-labeling assays, where Turbo-ID fused to the TTC5 mutants biotinylated tubulins less effectively than wild type TTC5 (**Figure S3G**).

In contrast to the interaction with αβ-tubulins, the TTC5 mutants were mostly competent for engagement of tubulin-synthesizing RNCs produced by *in vitro* translation. In this experiment, immunoprecipitation via tagged recombinant TTC5 followed by visualization of ribosomal proteins revealed that TTC5^D175A^ clearly engaged α- and β-tubulin RNCs, albeit somewhat less well than wild type TTC5 (**Figure 3F**). TTC5^T432E^ showed unimpaired engagement of β-tubulin RNCs but partial impairment for α-tubulin RNCs compared to wild type TTC5 (**Figure 3F**). Thus, these mutants are mostly selective in their inability to be sequestered by αβ-tubulins, while still retaining their capacity to engage tubulin-synthesizing ribosomes.

When introduced into TTC5 knockout cells, both TTC5^D175A^ and TTC5^T432E^ showed markedly reduced baseline tubulin mRNA levels despite unaffected transcription as evidenced by unchanged levels of unspliced pre-mRNAs (**Figure 3G, S3H**). These data are consistent with constitutive activation of the TTC5 mutant in the absence of efficient repression via αβ-tubulins. Interestingly, cells expressing these mutants still responded to colchicine treatment with further degradation of tubulin mRNAs (**Figure 3H, S3I**). This might be due to a weak but still relevant binding of the TTC5 mutants to αβ-tubulins that is further lost upon colchicine, or could hint at another colchicine-regulated factor such as SCAPER. This remains to be investigated. Regardless, the data illustrate that point mutants that relatively selectively impair the TTC5 interaction with αβ-tubulins result in loosened regulation of tubulin mRNA degradation. We therefore conclude that αβ-tubulins are physiologically relevant regulators of TTC5 activity.

### Constitutive activation of TTC5 causes mitotic defects

To investigate the physiological importance of αβ-tubulin-mediated repression of TTC5 we assessed mitotic fidelity in living cells using microscopy (**Figure 4A**). In agreement with previous reports, compared to parental cells, TTC5 knockout cells showed a higher rate of errors in chromosome alignment onto the metaphase plate (2.8-fold, **Figure 4B**), a higher rate of chromosome segregation errors in anaphase (2.6-fold, **Figure 4C**), and a subtle but highly reproducible delay in mitotic progression (**Figure S4B**). These phenotypes were rescued by re-expression of TTC5^WT^. Notably, reconstitution of TTC5 KO cells with the constitutively active TTC5^D175A^ and TTC5^T432E^ mutants showed similar types and frequencies of errors as in the knockout cells (**Figures 4B, 4C**, **S4A-C**). Thus, both the absence of TTC5-mediated autoregulation, which results in moderately higher tubulin mRNA than normal^15^, and constitutively active TTC5, which results in lower tubulin mRNA than normal, impair mitotic fidelity. These results underscore the importance of maintaining tubulin mRNA levels within a specific and narrow range via a fully functional dynamic autoregulation system.

**Figure 4.**
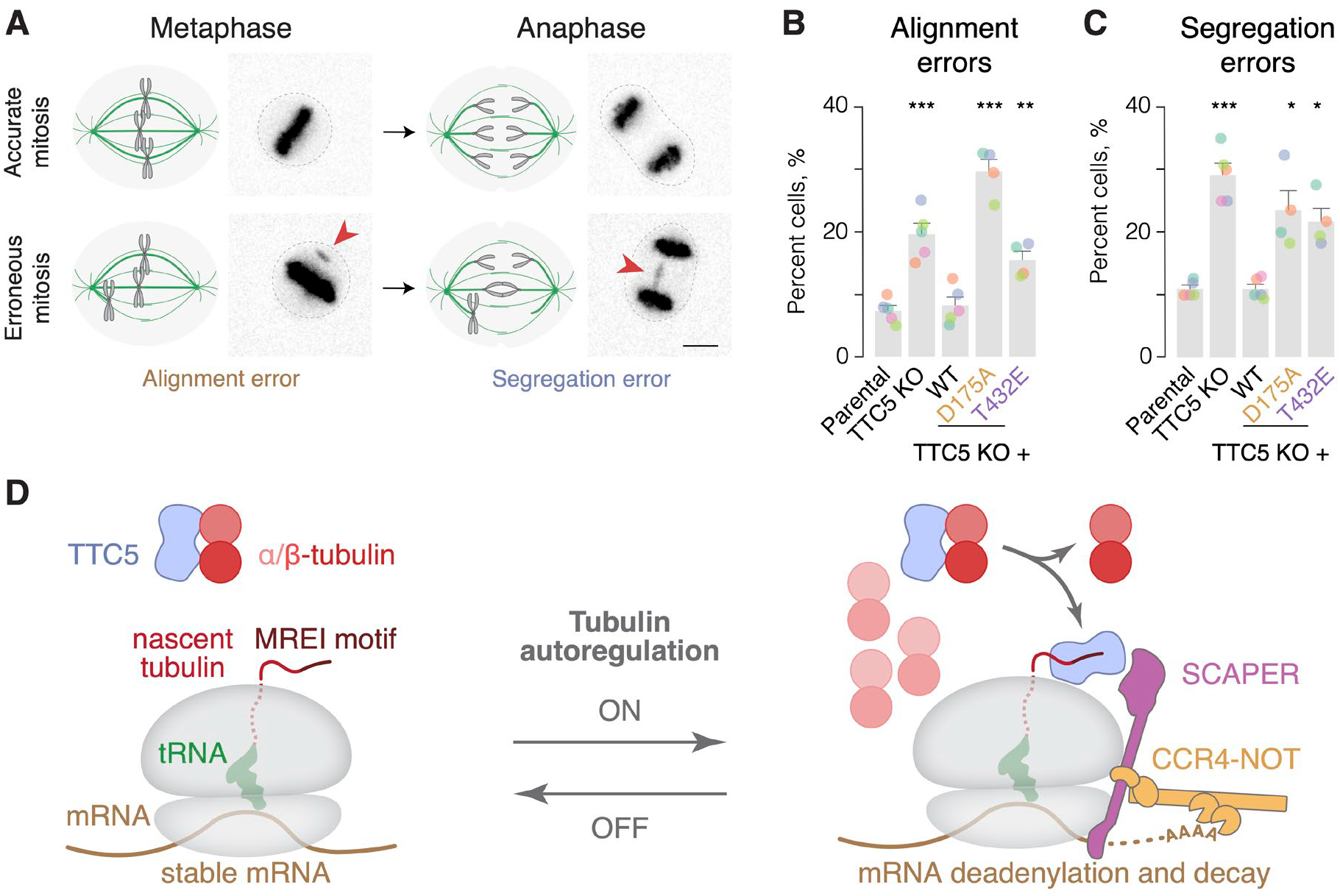
Constitutive activation of TTC5 causes mitotic defects. **(A)** Schematic and representative images of errors in chromosome alignment onto the metaphase plate and segregation in anaphase. Scale bar = 5 μm. (B-C) Occurrence of errors in chromosome alignment onto the metaphase plate **(B)** and errors in chromosome segregation in anaphase **(C)** in HeLa parental, TTC5 knockout, and the indicated Flag-TTC5 rescue cell lines (mean ± SD from at least three independent replicates represented with color-coded dots, and with at least 100 quantified cells per cell line). Single, double, and triple asterisks indicate p < 0.05, p < 0.01, and p < 0.001, respectively, in unpaired, two-tailed Student’s t tests for each of the indicated cell lines with the parental cell line as reference. **(D)** Proposed model for tubulin autoregulation.

## Discussion

Tubulin autoregulation was discovered over four decades ago, yet the control mechanisms governing it have long remained enigmatic. Early studies delineated the co-translational nature of this pathway and its sensitivity to microtubule depolymerizing drugs^9,11–13,31,32^. Recent investigations have elucidated the molecular players proximal to the ribosome, leading to a model where TTC5 engagement of nascent tubulin initiates a cascade involving SCAPER and the CCR4-NOT deadenylase complex, culminating in tubulin mRNA degradation^14,15^. Because this cascade is not constitutively active, one or more of its components must be under regulatory control to allow selective and dynamic activation of tubulin mRNA decay. These control point(s) have long been unclear, but are now amenable to study via TTC5, SCAPER, and CCR4-NOT.

Our study uncovers a crucial regulatory step in this pathway of selective mRNA decay: the reversible sequestration of TTC5 by soluble αβ-tubulins mediated by a previously overlooked C-terminal domain of TTC5. Under normal conditions, soluble αβ-tubulins sequester TTC5, preventing its engagement with RNCs (**Figure 4D**). Given a Kd of 0.5 μM between αβ-tubulin and TTC5, roughly 85% of TTC5 (∼35 nM in cells) would be sequestered by the ∼3 μM soluble αβ-tubulin. The remainder (∼5 nM) may be responsible for the low-level constitutive turnover of tubulin mRNA, explaining why TTC5 knockout cells have modestly elevated tubulin mRNA levels even in unperturbed conditions at steady state^15^.

Upon microtubule destabilization, αβ-tubulins progressively lose their capacity to interact with TTC5, allowing TTC5 to engage tubulin-translating ribosomes and trigger mRNA degradation (**Figure 4D**). Because the loss of interaction can be recapitulated using recombinant immobilized TTC5 in pulldown assays, we speculate that some change to αβ-tubulins triggered by microtubule depolymerization is responsible. Furthermore, the loss of interaction occurs progressively over 30-60 minutes and cannot be recapitulated with depolymerizing drugs in vitro. One explanation is a model where posttranslational modification(s) on αβ-tubulins progressively change, reducing their affinity to TTC5. Alternatively, or concomitantly, other proteins with higher affinity for soluble αβ-tubulins may become available and outcompete TTC5. Given the high complexity of the tubulin gene network^33^, vast tubulin posttranslational modifications^34^ and regulatory interaction partners^5^, the possibilities are myriad. An important future goal is to elucidate the mechanism that controls the interaction between TTC5 and αβ-tubulins.

The fact that TTC5’s C-terminal tail is involved in both its interaction with αβ-tubulins and with tubulin-synthesizing ribosomes is noteworthy. It helps explain why the two interactions seem to be mutually exclusive, and adds to our understanding of how the N-terminus of nascent tubulins is recognized. A combination of AlphaFold2 prediction and reinterpretation of earlier cryo-EM data support a role for the C-terminal tail in stabilizing the TTC5-RNC interaction. This structural model should be considered provisional until higher resolution structures are available. Nonetheless, the model explains why deletion of this tail impairs RNC recognition, and identifies this previously overlooked element of TTC5 as a crucial regulatory switch involved in both repression and activation of tubulin mRNA decay.

The ability to control TTC5 activity by αβ-tubulins is of physiological importance. TTC5 mutants that impair αβ-tubulin-mediated sequestration are constitutively active, leading to diminished tubulin mRNA levels and compromised chromosome alignment and segregation during mitosis. These same endpoint phenotypes were also seen in TTC5 and SCAPER knockouts and inactivating mutants, where autoregulation is completely lost and steady-state tubulin mRNA levels are higher than normal^14,15^. This is because mitotic fidelity is extraordinarily sensitive to altered microtubule dynamics and biomechanical properties of the mitotic spindle^35^. Both microtubule stabilization and destabilization cause similar loss of mitotic fidelity^23^. Errors in chromosome alignment and segregation commonly result in aneuploidy and DNA damage^36,37^, both of which have been linked to human disease, such as cancer^38–42^.

Mutations in TTC5 and SCAPER have also been implicated in neurodevelopmental disorders^16–18^, underscoring the broader importance of tubulin autoregulation in human physiology. Tubulin autoregulation is highly conserved amongst higher eukaryotes, operating in all cell types tested so far^24,43–47^. Yet for their proper functioning, differentiated cells require different ratios of soluble versus polymerized αβ-tubulins. Furthermore, this ratio may also need to be adjusted dynamically in response to physiologic context or different stages of the cell cycle. How cells fine-tune the tubulin autoregulation pathway to respond to context- and cell type-dependent needs remains to be elucidated. Our discovery of αβ-tubulins as a key control point for TTC5 regulation opens the door to elucidating the signals that regulate this step. Analogous regulatory mechanisms may operate on other steps in this pathway including the C-terminal molecular switch element in TTC5 and SCAPER, a known target of cell cycle regulated kinases^48–50^.

## Acknowledgements

We are grateful to the Proteins, peptides and RNA to protein core facility at the University of Geneva in Switzerland for their help with mass spectrometry; Vincent Mercier and the ACCESS platform for their help with high-throughput imaging and data analysis for the BiFC experiments; M. Skehel, S.-Y. Peak-Chew and C. Franco for help with mass spectrometry experiments and data handling; Michel Steinmetz and his team, Orsolya Barabas, Vladimir Arankin, Andreas Boland, Lina Poulain, and Ana Hoefler for their help and advice; all members of the Gasic and Lin teams for fruitful discussions and support.

This work was supported by the Swiss National Science Foundation Eccellenza Fellowship (SNSF PCEFP3_194312 to I.G.), the Republic and Canton of Geneva, Switzerland (DIP, to I.G.), National University of Singapore (PYP Start-up grant to Z.L.), Ministry of Education, Singapore (MOE AcRF Tier 1 A-8000057-00-00 to Z. L.), Medical Research Council, as part of United Kingdom Research and Innovation (MC_UP_A022_1007 to R.S.H.). A. B. is a recipient of the Salary Award from the Institute of Genetics and Genomics at the University of Geneva (iGE3). A.C.A. and M.H received funding from EMBO (EMBO ALTF 258-2023 to A.C.A, and ALTF 116-2020 to M.H.). M.H. received funding from the European Union’s Horizon 2020 research and innovation programme under the Marie Sklodowska-Curie grant agreement no 101029853. I.G. was the Dale F. Frey Breakthrough Scientist of the Damon Runyon Cancer Research Foundation (DRG:227916).

## Author contributions

A.B. and D.T.E.J co-discovered the C-terminal tail of TTC5 as a molecular switch in tubulin autoregulation, with structural insight from M.H.. A.B characterized αβ-tubulin/TTC5 complex, generated and tested all the TTC5 mutant cell lines, and performed most experiments (**Figures 2E-G**, **3A-B, 3H**, **S2D**, **S3A-B, S3G, S3I**). D.T.E.J. performed proximity labeling with TTC5 C-terminal truncation mutants of TTC5 (**Figure S2C**). M.H. performed *in vitro* translation assays, GFP-TTC5 and proteomic proximity labeling experiments (**Figures 1G, 2H**, **3F**, **S1H, S1J-K**). A.C.A. designed and generated the BiFC assays and performed the related experiments (**Figures 3C-E**, **S3C-F**). O.V. generated and analyzed HDX-MS data (**Figures 2A-D**, **S2A-B**). E.V. purified the recombinant TTC5 proteins and did western blots (**Figure S1A, S1I**, **S4C**). Z.L. discovered αβ-tubulins as repressors of TTC5 while he was a postdoctoral fellow with R.S.H (**Figures 1B-E**, **S1D-E, S1G**). I.G. designed and performed GFP-TTC5 abundance and localization experiments, AlphaFold2 predictions, some pulldowns, mRNA measurements, and phenotypic analysis of mitosis, (**Figures 1A, 1F**, **2I**, **3G**, **4A-D**, **S1A-C, S1F, S1I**, **S2E-G**, **S3H** and **S4A-B**). I.G., Z.L., and R.S.H., supervised different parts of the project. I.G. conceived the project with input from R.S.H. and M.H., oversaw its implementation, and wrote the manuscript. All authors contributed to manuscript editing.

## Materials and methods

### Plasmids and reagents

Tubulin constructs (human TUBB and TUBA1B) for *in vitro* translation in rabbit reticulocyte lysate (RRL) were cloned into pCDNA3.1. Constructs used for the expression and purification of WT or mutant 6XHis-Twin-Strep-TTC5 recombinant proteins were cloned into pET28a vector. Mutant E.coli tyrosyl-tRNA synthetase for incorporation of Bpa was expressed and purified from pET21 vector as previously described^51^. Bacillus stearothermophilus suppressor tRNA^Tyr^ sequence^52^ carrying T7 promoter sequence at 5’ and a BSTN1 restriction site at 3’ was cloned into pRSET. For BiFC experiments, VC-TUBA1B/TUBB-P2A-VN-TTC5 constructs were synthetized by Twist Bioscience (used Venus sequence from Addgene plasmid #105804^53^) and TTC5 mutants generated by site-directed mutagenesis. For generation of stable cell lines, TTC5 WT or mutants were sub-cloned into the pcDNA 5/FRT/TO vector.

### Cell culture

Flp-In T-REx HeLa cells (Invitrogen) were maintained in DMEM supplemented with 10% fetal bovine serum. Rescue cell lines with stable expression of N-terminally-tagged Flag- or Strep-TTC5 (WT and mutants) were generated from the TTC5 knockout cells using the Flp-In system (K650001 Invitrogen) according to manufacturer’s protocol. Expression of transgene was induced with 200 ng/mL doxycycline for 20-48 hours. Colchicine, nocodazole, and combretastatin A4 treatments were performed in standard media at indicated concentrations and time duration.

### Recombinant protein and tRNA purification

WT and mutant 6XHis-Twin-Strep-tagged TTC5 were purified from E. coli (BL21) cells. Briefly, cells were transformed with pET28a plasmid encoding WT or mutant TTC5 and grown at 37 °C in LB containing 50 μg/mL kanamycin. Induction was done with 0.2 mM IPTG at an A600 of 0.6 at 16 °C overnight. For Bpa incorporation at position 194 in TTC5 protein, cells were co-transformed with pET28a plasmid encoding the amber mutant TTC5 and the pEVOL-pBpF plasmid (Addgene plasmid #31190). Cells were grown at 37 °C in LB containing 50 μg/mL kanamycin and 25 μg/mL chloramphenicol and induced with 0.2 % L-arabinose at an A600 of 0.3 for 30 min followed by a second induction with 0.2 mM IPTG at an A600 of ∼0.6 at 16 °C overnight. Bacterial lysate was prepared by French press in 50 mL cold lysis buffer (500 mM NaCl, 20 mM imidazole, 1 mM PMSF, 1 mM TCEP and 50 mM HEPES pH 7.4) per 1 L of culture. Clarified bacterial lysates from a 1 L culture were bound to 1 mL column of HisPur Ni-NTA Spin column (ThermoFisher) by gravity flow. Columns were washed with ∼10 column volumes of lysis buffer and eluted with 250 mM imidazole in lysis buffer. The eluate was then bound to 1 mL column of Streptactin Sepharose® resin (IBA 2-1201-002). After extensive washing with 500 mM NaCl, 1 mM TCEP and 20 mM HEPES pH 7.4, the bound TTC5 protein was eluted with washing buffer containing 50 mM biotin and dialyzed against dialysis buffer (500 mM NaCl, 1 mM TCEP and 20 mM HEPES pH 7.4). E.coli Bpa tyrosyl-tRNA synthetase was purified via the C-terminal His tag on a Ni-NTA column, desalted by a gel filtration column on FPLC and concentrated by Amicon Ultra centrifugal filter (Millipore, Z717185-8EA). *B. stearothermophilus* tRNA^Tyr^, was synthesized by *in vitro* transcription. The pRSET-based construct was digested with BSTN1, yielding a DNA fragment containing the exact tRNA^Tyr^ sequence under a T7 promoter. 5 mL transcription reaction was carried out with 1.2 mg DNA template, 1 mM spermidine, 5 mM DTT, 0.1 % Triton, 5mM NTPs, 25 μM MgCl_2_, 20 μg/mL E. coli pyrophosphatase, 20 μg/mL T7 polymerase and 125 U Recombinant RNasin (Promega) for 4 hours at 37 °C. The reaction product was digested with Turbo DNase (Ambion) and extracted by acid phenol chloroform extraction to yield purified tRNA.

### Western blot

Protein samples were resolved using 12% Tris-Tricine or 10% Bis Tris based gels followed by transfer to 0.2 mm nitrocellulose membrane (Amersham Cytiva). Primary antibody incubations were performed for 1 h at room temperature or 4 °C overnight. Detection was done using HRP-conjugated secondary antibodies and SuperSignal West Pico Chemiluminescent substrate (Thermo Fisher), or DyLight conjugated antibodies (ThermoFisher) and Odyssey Infra-Red Imaging System (LI-COR). Expression of N-Flag WT or mutant TTC5 in rescue cell lines was detected by anti-Flag M2-HRP (Sigma A8592), monoclonal anti-Flag M2 (Sigma F3165), or anti-Strep antibody (abcam ab76949). RPL8 protein was detected by anti-RPL8 antibody (abcam ab169538), β-tubulin by antibodies (Cell Signaling Technologies 2128, Sigma-Aldrich T7816), α-tubulin by monoclonal antibody DM1A (Invitrogen 14-4502-37), and GAPDH with anti-GAPDH antibody (ThermoFisher MA515738). TTC5 was detected by using anti-TTC5 antibodies (Epigentek A66330, Novus Biologicals NBP1-76636, and ProSci 3053). BiFC constructs were detected using a rabbit polyclonal anti-GFP antibody (Torrey Pines Biolabs, TP401).

### *In vitro* transcription and translation

All *in vitro* transcription of tubulin constructs utilized PCR product as template. The 5’ primer contains the SP6 promoter sequence and anneals to the CMV promoter of pCDNA3.1. The 3’ primers anneal at codon 54-60 or 84-90 of nascent tubulin and contain extra sequence encoding MKLV to generate 64-mer or 94-mer constructs respectively. Transcription reactions were carried out with SP6 polymerase for 1 hour at 37 °C. Transcription reactions were directly used for *in vitro* translation in a homemade rabbit reticulocyte lysate (RRL)-based translation system as previously described^54^. For incorporation of Bpa by amber suppression, 5 μM B. Stearothermophilus tRNA^Tyr^, 0.25 μM Bpa tyrosyl-tRNA synthetase, and 0.1 mM Bpa were included in the translation reaction. As indicated in the figure legends, WT or mutant 6XHis-TwinStrep-tagged TTC5 were included in the translation reactions. Translation reactions were performed at 32 °C for 15–20 min. For analysis of total translation level of nascent chains, a 1uL aliquot of the translation reaction was mixed with protein sample buffer and analyzed by SDS-PAGE and autoradiography.

### *In vitro* analysis of TTC5 binding to tubulin RNCs

To test binding of WT or mutant TTC5 variants, 20–40 μl translation reaction of nascent α- or β-tubulin 64-mer containing 100–250 nM WT or mutant 6XHis-Twin-Strep-tagged TTC5 were carried out. The reactions were diluted 10 folds with PSB and incubated with 5 μL Streptactin Sepharose (IBA 2-1201-010) at 4 °C for 2 hours. Beads were washed four times with 400 μl PSB and eluted with 20 μL of 50 mM biotin in PSB at 4 °C for 30 min. Eluates were mixed with protein sample buffer for SDS-PAGE and analyzed with SYPRO Ruby protein gel stain (ThermoFisher S12000), autoradiography or western blotting.

### Autoregulation assay

Parental HeLa T-REx, TTC5 knockout and indicated rescue cell lines were grown to ∼70 % confluency in 6-well plates in the presence of 200 ng/ml doxycycline for 24 hours. To activate tubulin autoregulation pathway media containing either DMSO (vehicle control) or microtubules destabilizing drug colchicine (1 μM) was added to cells for 7 hours. Cells were harvested by scraping in RA1 lysis buffer and total RNA was isolated using the NucleoSpin RNA Mini Kit for RNA Isolation (Macherey-Nagel, 740955) according to manufacturer’s protocol. 1μg of total RNA was used to synthesize cDNA using the SensiFAST cDNA Synthesis Kit (Bioline, BIO-65054) following manufacturer’s instructions. qPCR was carried out using 10 ng of cDNA and 2x PowerUp SYBR Green master mix (Life Technologies, A25777) and indicated primers on a BioRad thermocycler (BioRad). Following primers were used for qPCR:

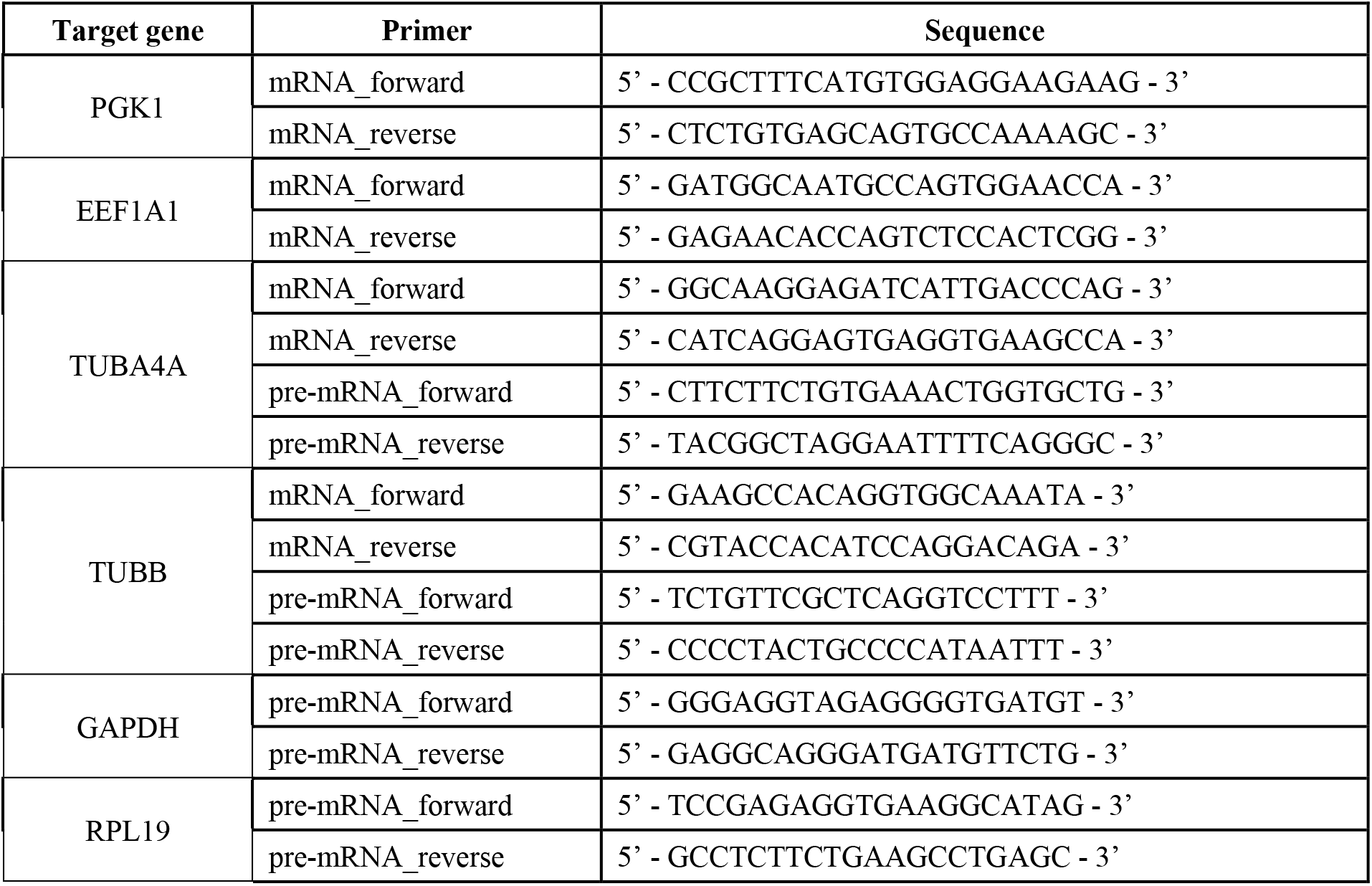

Data analysis was performed using the ddCt method^55^. All data were normalized to reference housekeeping genes, and to either DMSO treated controls or parental cell line as indicated. Experiments include at least three biological replicates. Processing, statistical analysis, and data plotting were performed in R.

### Recombinant TTC5 pulldown assays of cell lysates. T

TC5 knockout cells were grown to 70-80 % confluency in 145 mm dish and treated with DMSO control, colchicine (10 μM) or nocodazole (10 μM) for the times indicated in the figure legends. For preparation of cytosolic cell lysates, cells were pelleted by centrifugation at 500 g for 5 min and lysed with lysis buffer (100 mM KAc, 5 mM MgAc2, 1 mM DTT, 100 μg/mL digitonin, 1X EDTA-free protease inhibitor cocktail (Roche) and 50 mM HEPES pH 7.4) for 10 min on ice. Lysates were cleared by centrifugation at 20000 g for 15 min at 4 °C. Lysate concentrations were determined by Pierce BCA assay kit (ThermoFisher). An aliquot of the lysates was used for total RNA extraction and analyzed for tubulins mRNA by RT-qPCR as described above. Another aliquot of lysate was incubated with 500 nM recombinant TTC5 for 2 min on ice followed by incubation with 10 μl Streptactin Sepharose for 2 hours at 4 °C to recover TTC5 and all bound components. Control samples omitted recombinant TTC5. Beads were washed three times with 400 μl PSB and eluted with 50 μL of 50 mM biotin in PSB at 4 °C for 30 min. Eluted samples were analyzed by SDS-PAGE and SYPRO Ruby stain to visualize TTC5 and associated proteins.

### Flag- and Strep-TTC5 pulldowns from stable cell lines

To analyze interaction partners of TTC5^WT^ and indicated mutants we used HeLa T-REx TTC5 KO cells expressing either Flag- or Twin-Strep-tagged version of TTC5 under the doxycycline-inducible promoter. Cells were grown in 150 mm plates up to 80-90 % confluency in the presence of 200 ng/ml doxycycline for at least 20 hours. Cells were washed once with ice-cold PBS, pelleted, and cytosolic extracts were prepared by lysis in 1 ml digitonin lysis buffer for 10 min on ice (50 mM HEPES pH 7.4, 100 mM KAc, 5 mM MgAc_2_, 1 mM DTT, 1x EDTA-free protease inhibitor cocktail (Roche), 100 μg/ml digitonin). Lysates were cleared by centrifugation at maximum speed at 4 °C for 15 min, followed by 1 hour incubation at 4 °C with 20 μl of either Anti-FLAG M2 Magnetic Beads (Sigma M8823) for Flag-TTC5 cell lines, or MagStrep Strep-Tactin beads (IBA 2-1613) for Twin-Strep-TTC5 cell lines. Beads were then washed four times with physiological salt buffer (PSB: 50 mM HEPES pH 7.4, 100 mM KAc, 2 mM MgCl2, 10 μg/ml digitonin) with a change of tube for the last wash. Elution was done by adding either 20 μl of 0.2 mg/ml of 3xFLAG peptide (Sigma-Aldrich F4799) or 20 μl of 1x Strep-Tactin Elution Buffer (IBA 2-1042-025) to the beads at 4 °C for 30 min. Eluted proteins were separated by SDS-PAGE and analyzed by western blot with indicated antibodies.

### Microtubule co-pelleting assay

5 % glycerol and 1 mM GTP were added to stock porcine tubulin (7 mg/mL) in General Tubulin Buffer (GTB: 2 mM MgCl2, 0.5 mM EGTA, 80 mM PIPES pH 6.9) and incubated at 37°C for 20 min to assemble microtubules in vitro. 20 μM taxol was added to stabilize polymerised microtubules after incubation. Recombinant TTC5 was buffered-exchanged into binding buffer (100 mM KCl, 5 mM MgCl2, 1 mM EGTA and 20 mM HEPES pH 7.4). 2.5 μM TTC5 was incubated with 0.6 mg/mL assembled microtubules at room temperature for 30 min. A 50 μl mixture was layered on a 100 μl glycerol cushion (100 mM KCl, 5 mM MgCl2, 1 mM EGTA, 60 % Glycerol, 20 μM taxol and 20 mM HEPES pH 7.4) and centrifuged at 100000 g at room temperature for 40 min to pellet the microtubules. The supernatant was carefully removed, and the pellet was resuspended in GTB. Proteins in supernatant and resuspended pellet were analyzed by SDS-PAGE followed by SYPRO Ruby staining.

### Measurement of TTC5 and tubulin binding constant

20 μM recombinant TTC5 was labelled with 22 μM Oregon GreenTM 488 Maleimide (ThermoFisher O6034) in 500 mM NaCl, 1 mM TCEP and 20 mM HEPES pH 7.4 and incubated on ice for 30 min. Excess dye was removed by desalting column. 50 nM labelled TTC5 was mixed with various porcine tubulin concentrations in 250 mM NaCl, 1 mM TCEP and 20 mM HEPES pH 7.4 and measured with microscale thermophoresis (2bind) to generate a binding curve. Measurements were performed in triplicates.

### Bimolecular fluorescence complementation (BiFC) analysis

At day 0, 18000 Flp-In T-REx HeLa cells of the indicated genotypes were seeded in an μ-Plate 96-well plate (ibidi #89626) in the presence of 200 ng/mL doxycycline. After 6 hours, medium was exchanged to DMEM without phenol-red (Thermo Fisher Scientific #21063029) supplemented with 10 % fetal bovine serum, 50 nM SiR-DNA (Spirochrome #SC007) and 200 ng/mL doxycycline. Cells were incubated for 16 hours prior to imaging. High-throughput live cell imaging was performed in an ImageXpress Micro Confocal automated microscope (Molecular Devices™, wide-field mode) equipped with a 40x water immersion objective (0.95 NA, Nikon). A total of 8-12 regions of interest were acquired per well. Image segmentation was performed using a custom module editor MetaXpress from Molecular Devices, by generating masks to extract Venus’s fluorescence intensity across all conditions. Briefly, cell nuclei and body masks were created using SiR-DNA to define a master object (all cells). Venus-positive cells were identified based on average fluorescence intensity and values plotted using GraphPrism 8.

### Proximity labeling in **Fig. S1J** and **S1K**

HEK T-REx TTC5 KO cells were complemented with TurboID-FLAG-TTC5 constructs expressed from stably integrated pcDNA5/FRT/TO plasmid. For each condition, a 145 mm plate of WT cells or TurboID-TTC5 cells was grown to 70% confluency. To avoid strong overexpression of TurboID-TTC5, leaky expression from the doxycycline-inducible promoter was used without addition of doxycycline. Cells were treated for 30 min with 10 μM colchicine as indicated and then 50 μM biotin (APExBIO A8010) was added for another 2.5 hours. Cells were washed once with ice-cold PBS, pelleted, and cytosolic extracts were prepared by lysis in 0.8 ml digitonin lysis buffer per plate for 10 min on ice (50 mM HEPES pH 7.4, 100 mM KAc, 5 mM MgAc_2_, 1 mM DTT, 1x EDTA-free protease inhibitor cocktail (Roche), 0.01% digitonin). Lysates were cleared by centrifugation at 20000 g at 4 °C. Lysates were then incubated on a rotating wheel with 20 μl of streptavidin-coupled magnetic beads (Pierce 88817) for 1.5 hours at 4 °C. Beads were then washed with 1 ml each of physiological salt buffer (PSB: 50 mM HEPES pH7.4, 100 mM KAc, 2 mM MgAc_2_) with 0.01 % digitonin, wash buffer 1 (1 % SDS, 10 mM Tris-HCl pH 8), wash buffer 2 (1 M NaCl, 10 mM Tris-HCl pH 8, 0.01% digitonin), and wash buffer 3 (2 M urea, 10 mM Tris-HCl pH 8, 0.01% digitonin), and PSB with 0.01% digitonin. Beads were transferred to a fresh tube with the last wash and eluted with 20 μl sample buffer supplemented with 2 mM biotin for 5 minutes at 95 °C. Eluted proteins were separated by SDS-PAGE and analyzed by total protein staining with SYPRO Ruby, or by western blot. For mass spectrometry comparison of biotinylated proteins under control (DMSO) and colchicine-treated conditions, we reanalyzed a previously published dataset of TurboID-TTC5 proteomics data^15^. Samples were prepared as described above with minor modifications and quantified with tandem mass tag-labelling of peptides followed by proteomic analysis as described previously^15^.

### Proximity labeling in **Fig. S3G**

TurboID-Flag construct was fused to the N-terminus of TTC5 (WT or mutants) and cloned into pcDNA5/FRT/TO vector. HeLa T-REx TTC5 KO cell line was used to create rescue (WT) or indicated mutant cell lines with stable expression of TurboID-Flag-TTC5. For western blot of biotinylated proteins cells were seeded at 70 % confluency and induced with 5 ng/ml doxycycline for 24 hours. 50 μM biotin was added to cells for 15 min at 37 °C followed by five washes with ice-cold PBS and cell lysis with RIPA buffer (50 mM Tris HCl pH 8, 150 mM NaCl, 1 % Triton X-100, 0.5 % sodium deoxycholate, 0.1 % SDS). Total protein concentration was measured with Pierce BCA Protein Assay Kit (Thermo Scientific 23227) and 15 μg of total protein lysates was analyzed by western blot. Biotynylated proteins were visualized with Streptavidin antibody conjugated to horseradish peroxidase (Invitrogen S911). Experiment was performed in four biological replicates. Quantification of tubulin biotinylation was done using Fiji. The signal intensity of biotinylated tubulin and TurboID-TTC5 bands were measured to calculate tubulin/TTC5-TurboID ratio for each cell line. TTC5 knockout cells were used as background control and TurboID-TTC5 WT sample served as a reference (100 % of biotinylated tubulin) to normalize data of TTC5 mutants.

### Analysis of TTC5 interaction partners by proteomics

To analyze in vivo interaction partners of TTC5 before and after colchicine treatment, we used 293 T-REx TTC5 KO cells complemented with doxycycline-inducible EGFP-tagged TTC5 as previously described^15^. Cells were grown in 145 mm plates to around 80 % confluence and treated with 10 μM colchicine for 0, 15, 30, 60, 120, or 180 minutes. Cells were washed once in ice-cold PBS, pelleted, and cytosolic extracts were prepared by lysis in 1 ml digitonin lysis buffer per plate for 10 min on ice (50 mM HEPES pH 7.4, 100 mM KAc, 5 mM MgAc_2_, 1 mM DTT, 1x EDTA-free protease inhibitor cocktail (Roche), 0.01% digitonin). Lysates were cleared by centrifugation at maximum speed at 4 °C in a table-top centrifuge. Lysates were then incubated on a rotating wheel with 10 μl of GFP-trap agarose (ChromoTek) for 1 hour at 4 °C. Beads were then washed twice with 1 ml each of physiological salt buffer containing 0.01 % digitonin, and twice with physiological salt buffer without detergent. For each timepoint, three technical replicates were subjected to label-free quantification by LC-MS/MS. The bead samples were buffer exchanged twice with 100 mM ammonium bicarbonate and on the last wash beads were left with minimum buffer to cover. The cysteines were reduced by adding 30 μL of 10 mM DTT and then alkylated with 30 uL of 55 mM iodoacetamide. Proteins were digested on beads with 1 μg trypsin (Promega, UK) for 18 hours at 37 °C. Peptides were acidified with the addition of 4 μl formic acid 2 % (v/v). The bead/peptide mix was then centrifuged at 14,000 x g for 5 minutes and the 20 μl of supernatant placed into a vial for LC-MS/MS analysis. LC-MS/MS was performed on an Ultimate U3000 HPLC (ThermoFisher Scientific, San Jose, USA) hyphenated to an Orbitrap QExactive Classic mass spectrometer (ThermoFisher Scientific, San Jose, USA). Peptides were trapped on a C18 Acclaim PepMap 100 (5 μm, 300 μm x 5 mm) trap column (ThermoFisher Scientific, San Jose, USA) and eluted onto a C18 Acclaim PepMap100 3 μm, 75 μm x 250 mm (ThermoScientific Dionex, San Jose, USA) using 30 minutes gradient of acetonitrile (4 to 30 %). For data dependent acquisition, MS1 scans were acquired at a resolution of 35,000 (AGC target of 1e6 ions with a maximum injection time of 50ms) followed by ten MS2 scans acquired at a resolution of 17,500 (AGC target of 2e5 ions with a maximum injection time of 100 ms) using a collision induced dissociation energy of 25. Dynamic exclusion of fragmented m/z values was set to 30 s. Raw data were imported and processed in MASCOT (Matrix Science). The raw files were submitted to a database search against the UniProt/SwissProt database. Database search parameters were set with a precursor tolerance of 10 ppm and a fragment ion mass tolerance of 0.8 Da. One missed enzyme cleavage was allowed and variable modifications for oxidized methionine, carbamidomethyl and phospho STY were included. The acquired LC-MS/MS raw files were processed using MaxQuant^56^ with the integrated Andromeda search engine (v1.6.6.0), and searched against Human Reviewed UniProt Fasta database. The MaxQuant output file (proteinGroups.txt) was then processed with Perseus software^57^. After uploading the matrix, the data was filtered to remove identifications from reverse database, identifications with modified peptide only, and common contaminants. Data were log2-transformed, a valid value filter was applied and missing values for remaining proteins were imputed with standard settings. We found that the 0- and 15-minute timepoints showed very similar interaction profiles and were thus grouped as “untreated” samples, and the 60-, 120-, and 180-minute timepoints showed highly similar profiles and were grouped as “colchicine-treated” samples for further analysis. To generate a volcano plot we compared all samples in the two groups. The 30-minute timepoint showed an intermediate profile and was omitted for this analysis.

### Hydrogen-deuterium exchange mass spectrometry (HDX-MS)

HDX-MS was performed at the UniGe Protein Platform (University of Geneva, Switzerland) following a well-established protocol with minimal modifications^58^. Details of reaction conditions:

| **Dataset :** | **State 1** | **State 2** |
| --- | --- | --- |
| *Description :* | TTC5 | TTC5 + abTubulin |
| Reaction volume : | 50 ul | 50 ul |
| % D2O in the reaction : | 89.0% | 89.0% |
| temperature : | 0 °C | 0 °C |
| D2O incubation times (sec) : | 3s, 30s, 300s | 3s, 30s, 300s, |
| Control sample : | Non-deuterated (ND) TTC5 |  |
| Quench buffer : | 4M Gdn-HCl/ 1M NaCl/ 100mM NaH<sub>2</sub>PO<sub>4</sub> pH 2.4/ 1% FA |  |
| Quench buffer volume : | 20 ul | 20 ul |
| Number of peptides analyzed : | 121 | 121 |
| Sequence coverage : | 99.0% | 99.0% |
| Replicates : | 3, 3, 3, 0, 0, 0 | 3, 3, 3, 0, 0, 0 |
| Criteria for HDX rate difference : | Difference of HDX level at a given timepoint is > 8 % and > 1 Da and p values of student t-test is < 0.01 |  |
| Protein amount (pmol) : | 78 | 78 + 116 |

HDX reactions were done in 50 μl volumes with a final protein concentration of 1.6 μM TTC5 and a 1.5-fold molar excess of αβ-tubulins. Briefly, 78 picomoles of protein were pre-incubated with αβ-tubulins or buffer for 1 h on ice in a final volume of 3 μl before deuteration. Deuterium on-exchange reaction was initiated by adding 47 μl of D2O exchange buffer (20 mM HEPES pH 7.5 / 150 mM NaCl / 1 mM TCEP in D2O) to the protein-ligand mixture. Reactions were carried-out on ice for three incubation times (3 s, 30 s, 300 s) and terminated by the sequential addition of 20 μl of ice-cold quench buffer (4 M Gdn-HCl / 1 M NaCl / 0.1 M NaH_2_PO_4_ pH 2.5 / 1 % Formic Acid). Samples were immediately frozen in liquid nitrogen and stored at -80°C for up to two weeks. All experiments were repeated in triplicates. To quantify deuterium uptake into the protein, samples were thawed and injected in a UPLC system immersed in ice with 0.1 % FA as liquid phase. The protein was digested via two immobilized pepsin columns (Thermo #23131), and peptides were collected onto a VanGuard precolumn trap (Waters). The trap was subsequently eluted, and peptides separated with a C18, 300 Å, 1.7 μm particle size Fortis Bio 100 x 2.1 mm column over a gradient of 8 – 30 % buffer C over 20 min at 150 ml/ min (Buffer B: 0.1 % formic acid; buffer C: 100 % acetonitrile). Mass spectra were acquired on an Orbitrap Velos Pro (Thermo), for ions from 400 to 2200 m/z using an electrospray ionization source operated at 300 °C, 5 kV of ion spray voltage. Peptides were identified by data-dependent acquisition of a non-deuterated sample after MS/ MS and data were analyzed by Mascot. Deuterium incorporation levels were quantified using HD examiner software version 3.3 (Sierra Analytics), and quality of every peptide was checked manually. Results are presented as percentage of maximal deuteration compared to theoretical maximal deuteration level. Changes in deuteration level between two states were considered significant if >12% and >1.4 Da and p< 0.01 (unpaired t-test) for a single deuteration time.

### Multiple sequence alignment (MSA) analysis

MSA of TTC5 protein was done using the ConSurf server^59^ (server for the identification of functional regions in proteins, https://consurf.tau.ac.il/consurf_index.php). Alignment was performed for AlphaFold2 predicted model of human TTC5 (AF-Q8N0Z6-F1). Exposed and buried residues of TTC5 were predicted with NACSES algorithm of ConSurf.

### Live cell imaging and data analysis

Flp-In T-REx HeLa cells of the genotypes indicated in the figure legends were plated in 8-well Lab Tek II Chamber 1.5 German coverglass dishes (Thermo Fisher 155409) in regular growth medium, and incubated for 6 hours. Medium was then changed to Leibovitz’s L-15 without phenol-red (ThermoFisher, 21083027) supplemented with 10% fetal bovine serum, 200 ng/mL doxycycline and 50 nM SiR-DNA (Cytoskeleton CY-SC007). Cells were incubated for 24 hours prior to imaging. Time lapse images were acquired using Nikon Eclipse Ti2-E inverted microscope (Nikon), equipped with Kinetix sCMOS camera (Photometrics), Spectrax Chroma light engine for fluorescence illumination (Lumencor), a perfect focus system, and an incubation chamber with 37 °C and controlled humidity (OkoLab). Three-dimensional images at multiple stage positions were acquired in steps of 2 μm, every 7 minutes for 10 hours using NIS Elements (Nikon) and 20x Plan Apochromat Lambda objective (NA 0.80, Nikon). Maximum intensity projections and inverted color profiles of representative examples of mitoses were prepared in Fiji and exported as still images. Analysis of mitotic cells was performed using 3D reconstructions in Fiji. The parameters scored (based on the SiR-DNA signal) were the occurrence of unaligned chromosomes in metaphase, and chromosome segregation errors in anaphase. Analyses of at least 100 cells per cell line in three biological replicates were documented using Excel and processed and plotted using R. Statistical analyses were performed in R. Instances where not all the chromosomes were properly aligned on the spindle equator in metaphase and/or anaphase are classified as chromosome alignment errors. Instances where sister chromatids failed to properly separate, either segregating both into the same daughter cell or forming a bridge in anaphase were classified as segregation errors. Numbers reported represent percentage of cells experiencing either abnormality.

## Supplementary figures, figure titles and legends

**Figure S1.**
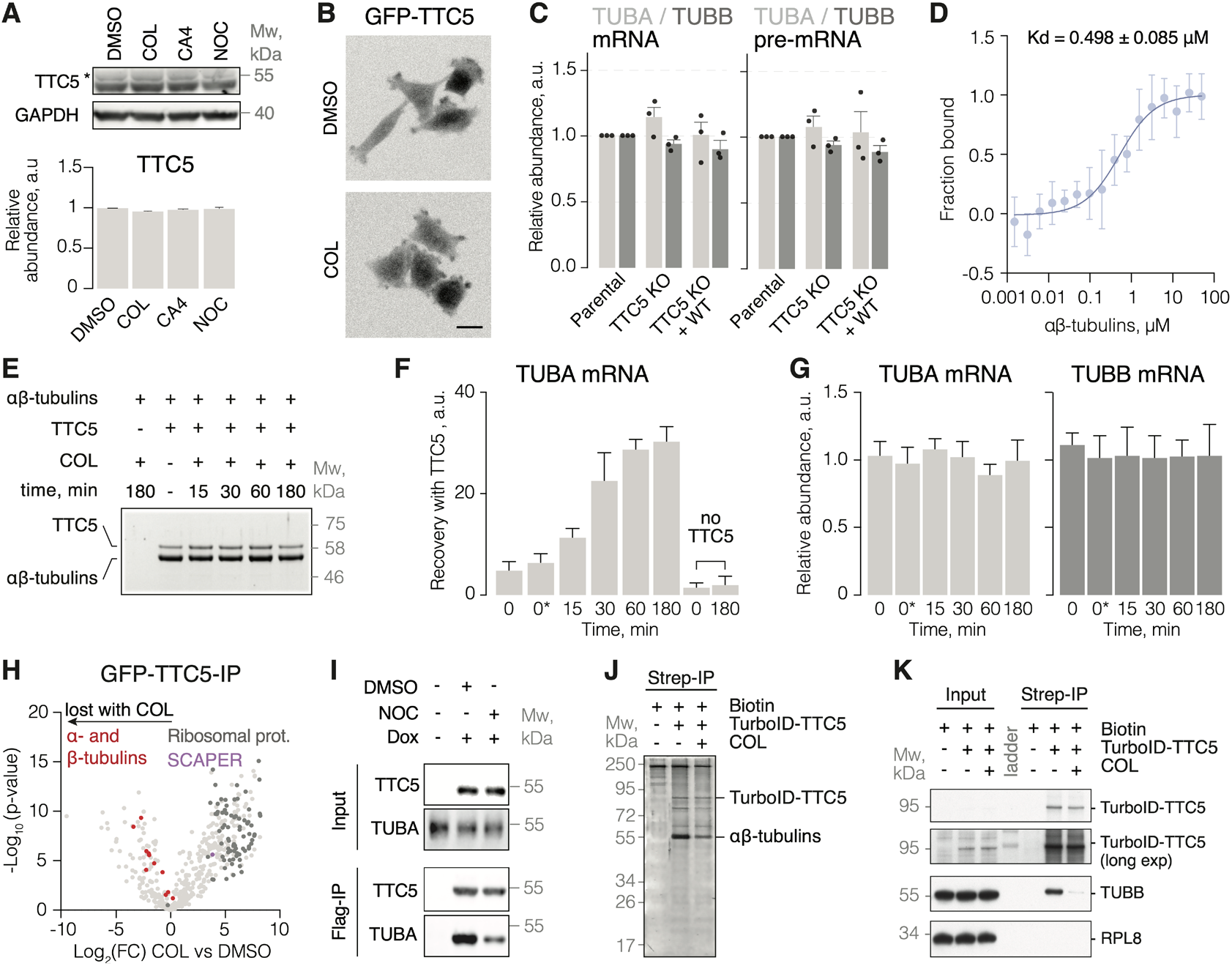
Soluble αβ-tubulins reversibly repress TTC5 to regulate its activity (related to **Figure 1**). **(A)** Total cell lysates from HeLa cells treated with DMSO or 1 μM colchicine (COL), combretastatin A4 (CA4) or nocodazole (NOC) for 5 hours. Proteins were separated by SDS-PAGE and visualized by western blot. TTC5 levels across the indicated conditions were normalized to GAPDH (mean ± SD from three independent replicates). Asterisk indicates TTC5 band. **(B)** Representative images of HeLa TTC5 knockout cells re-expressing GFP-TTC5, after treatment with DMSO control or 1 μM colchicine (COL) for 5 hours. Scale bar = 10 μm. **(C)** Relative α- and β-tubulin mRNA and pre-mRNA levels in HeLa parental, TTC5 knockout, and Strep-TTC5^WT^ rescue cell lines, normalized to a housekeeping transcript and the parental cell line (mean ± SD from three independent experiments). No statistically significant differences were detected in unpaired, two-tailed Student’s t-tests. **(D)** Dissociation constant of porcine tubulin and TTC5 determined by microscale thermophoresis. **(E)** Recombinant Strep-TTC5 was incubated with porcine brain tubulin in the absence or presence of 1 μM colchicine and the indicated time durations, then affinity purified via the Strep-tag. Pulled down proteins were separated by SDS-PAGE and visualized using western blot, showing that binding of TTC5 to purified porcine brain tubulin is not affected by prolonged incubation with colchicine in vitro. **(F)** The products recovered from cell lysates by binding to recombinant TTC5 shown in **Figure 1E** were analyzed for α-tubulin mRNA by quantitative RT-PCR (mean ± SD from three independent replicates). **(G)** Total α- and β-tubulin mRNA levels across the indicated conditions (mean ± SD from three independent replicates). **(H)** Total cell lysates of HEK293 TTC5 knockout cells re-expressing GFP-TTC5 and treated with DMSO control or 10 μM colchicine for 3 hours were immunoprecipitated using the GFP-tag. TTC5 interactome was subsequently analyzed using label-free quantitative mass spectrometry. Data are presented as Log2 fold change (Log2(FC)) in colchicine versus DMSO treated samples. Highlighted are the TTC5 interaction partners specifically lost or gained upon colchicine treatment. **(I)** HeLa TTC5 knockout cells re-expressing Flag-TTC5 under a doxycycline-inducible promoter were treated with DMSO or 1 μM nocodazole (NOC) for 3 hours. Cells were then lysed and TTC5 was affinity purified via the Flag-tag. Coimmunoprecipitated interactors were separated using SDS-PAGE and visualized using western blot. (J-K) HEK 293 TTC5 knockout cells re-expressing TTC5 N-terminally fused with TurboID biotin ligase under doxycycline-inducible promoter were treated with DMSO or 10 μM colchicine and incubated in the presence of 50 μM biotin for 2.5 hours. Biotynylated proteins were affinity purified, separated by SDS-PAGE and visualized using SYPRO Ruby stain **(J)** or western blot **(K)**. No statistically significant differences were detected in unpaired, two-tailed Student’s t-tests.

**Figure S2.**
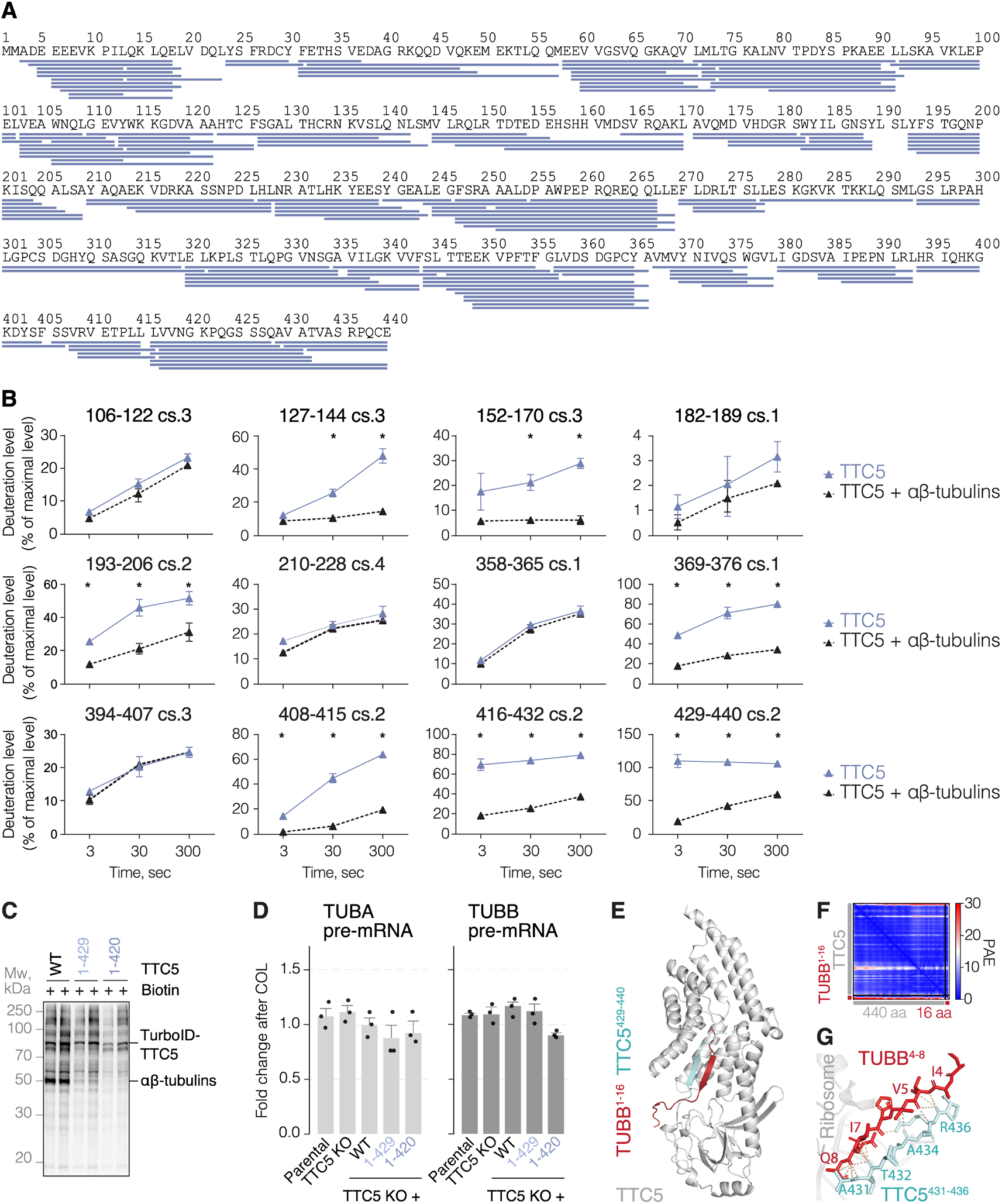
C-terminal tail mediates interaction with mature and nascent tubulin (related to **Figure 2**). **(A)** Peptide coverage for recombinant Strep-TTC5 related to the HDX-MS analysis presented in **Figure 2A-D**. **(B)** Deuteration profiles of a selection of TTC5 peptides upon incubation in deuterated water for the indicated time durations (mean ± SD from three independent replicates). **(C)** HeLa TTC5 knockout cells re-expressing TTC5 N-terminally fused with TurboID biotin ligase under doxycycline-inducible promoter were incubated in the presence of 50 μM biotin for 2.5 hours. Biotinylated proteins were separated by SDS-PAGE and visualized using western blot. Data from two biological replicates are shown. **(D)** Autoregulation assay with HeLa parental, TTC5 knockout, and the indicated Flag-TTC5 rescue cell lines. Data show the mean ± SD pre-mRNA levels after colchicine treatment, from at least three independent experiments. No statistically significant differences were detected in unpaired, two-tailed Student’s t-tests for each of the indicated cell lines with the DMSO-treated sample as reference. **(E)** AlphaFold2 multimer prediction of TTC5 (gray) bound to nascent β-tubulin (red). C-terminal tail of TTC5 (cyan) is predicted to form an anti-parallel beta sheet with nascent β-tubulin (red). **(F)** Predicted aligned error (PAE) for the predicted structure in panel E. **(G)** Close-up view of the C-terminal tail of TTC5 (cyan) and nascent β-tubulin (red) forming a beta sheet. Yellow dashed lines indicate possible interactions between TTC5 (cyan) and nascent β-tubulin (red).

**Figure S3.**
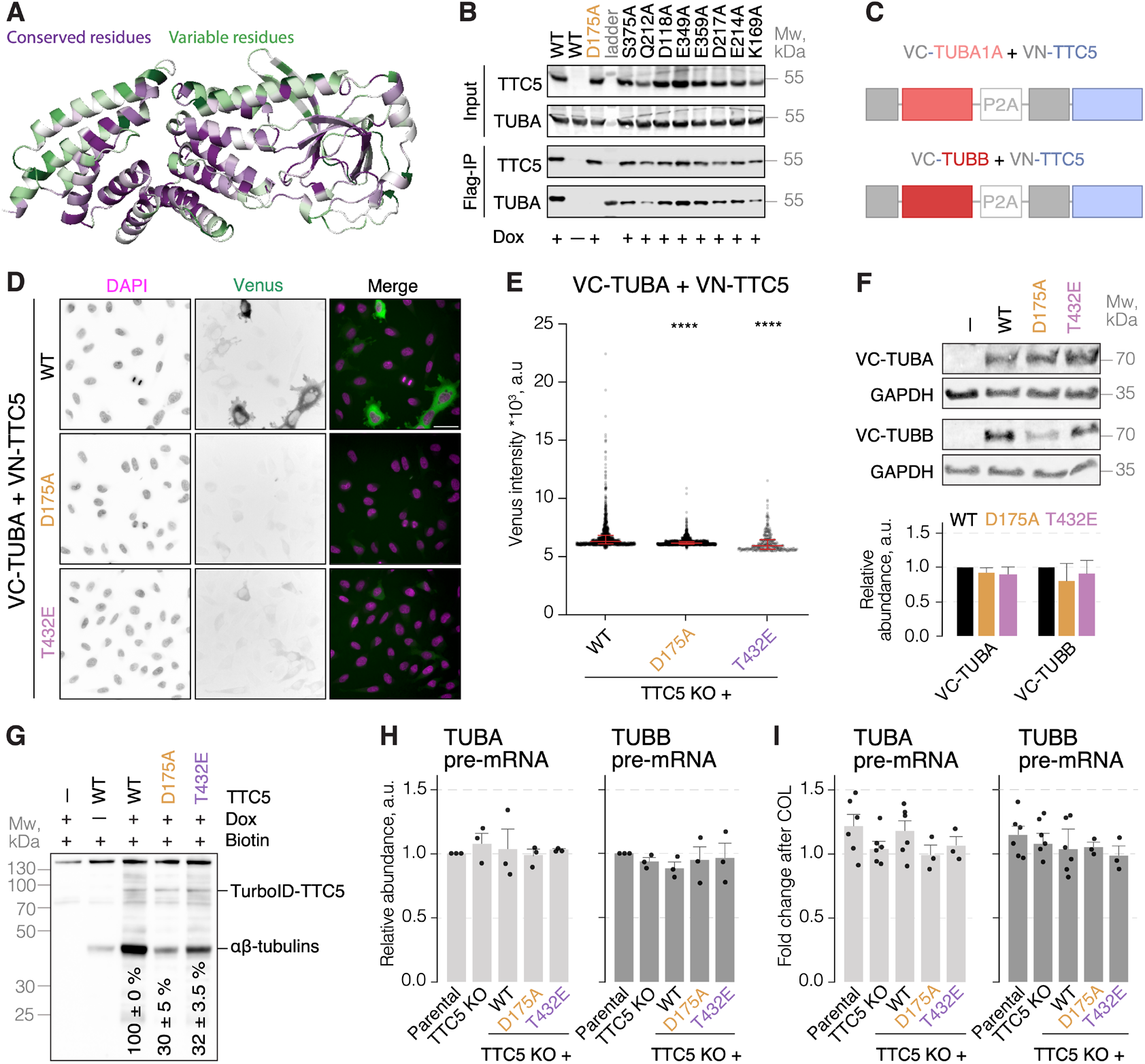
Loss of binding to αβ-tubulins constitutively activates TTC5 (related to **Figure 3**). **(A)** Multiple sequence alignment mapped onto the AlphaFold2-predicted structure of the human TTC5. **(B)** Indicated Flag-TTC5 constructs were expressed in HeLa TTC5 knockout cells under a doxycycline-inducible promoter and affinity purified via the Flag-tag. Coimmunoprecipitated interactors were separated using SDS-PAGE and visualized using western blot. **(C)** Schematic representation of the BiFC constructs. **(D)** Representative live-cell images of the indicated VN-TTC5 and VC-TUBA constructs expressed in HeLa TTC5 knockout cells. Scale bar = 20 μm. **(E)** Fluorescence intensity of Venus in cell lines expressing the indicated BiFC constructs. Dots represent measurements from individual cells, red line the median fluorescence intensity, and error bars interquartile range. Quadruple asterisks indicate p < 0.0001 in Mann-Whitney test for each of the indicated BiFC constructs with the BiFC constructs based on TTC5^WT^ as reference. **(F)** Total protein analysis by western blot for the indicated BiFC constructs in the cell lines used in panels D and E, and in **Figure 3D** and **3E**. Graph shows relative abundance of BiFC constructs normalized to housekeeping protein and TTC5^WT^ (mean ± SD from three independent experiments). No statistically significant differences were detected using unpaired, two-tailed Student’s t-test. Note that the polyclonal anti-GFP antibody only recognizes the VC fragment of Venus. **(G)** HeLa TTC5 knockout cells re-expressing TTC5 N-terminally fused with TurboID biotin ligase under doxycycline-inducible promoter were incubated in the presence of 50 μM biotin for 15 minutes. Biotinylated proteins were separated by SDS-PAGE and visualized using western blot. Numbers indicate remaining tubulin labeling normalized to the level of the relevant TurboID-TTC5 self-labeling and relative to tubulin biotinylation in cells expressing TurboID-TTC5^WT^. **(H)** Total pre-mRNA levels for α- and β-tubulin in the indicated HeLa cell lines. All the data were normalized to a reference transcript and the parental cell line (mean ± SD from three independent experiments). No statistically significant differences were detected using unpaired, two-tailed Student’s t-test. **(I)** Autoregulation assay with HeLa parental, TTC5 knockout, and the indicated Strep-TTC5 rescue cell lines. Data show pre-mRNA levels for α- and β-tubulin normalized to a reference transcript and the DMSO vehicle-treated sample for each tested cell line (mean ± SD from at least three independent experiments). No statistically significant differences were detected using unpaired, two-tailed Student’s t-test.

**Figure S4.**
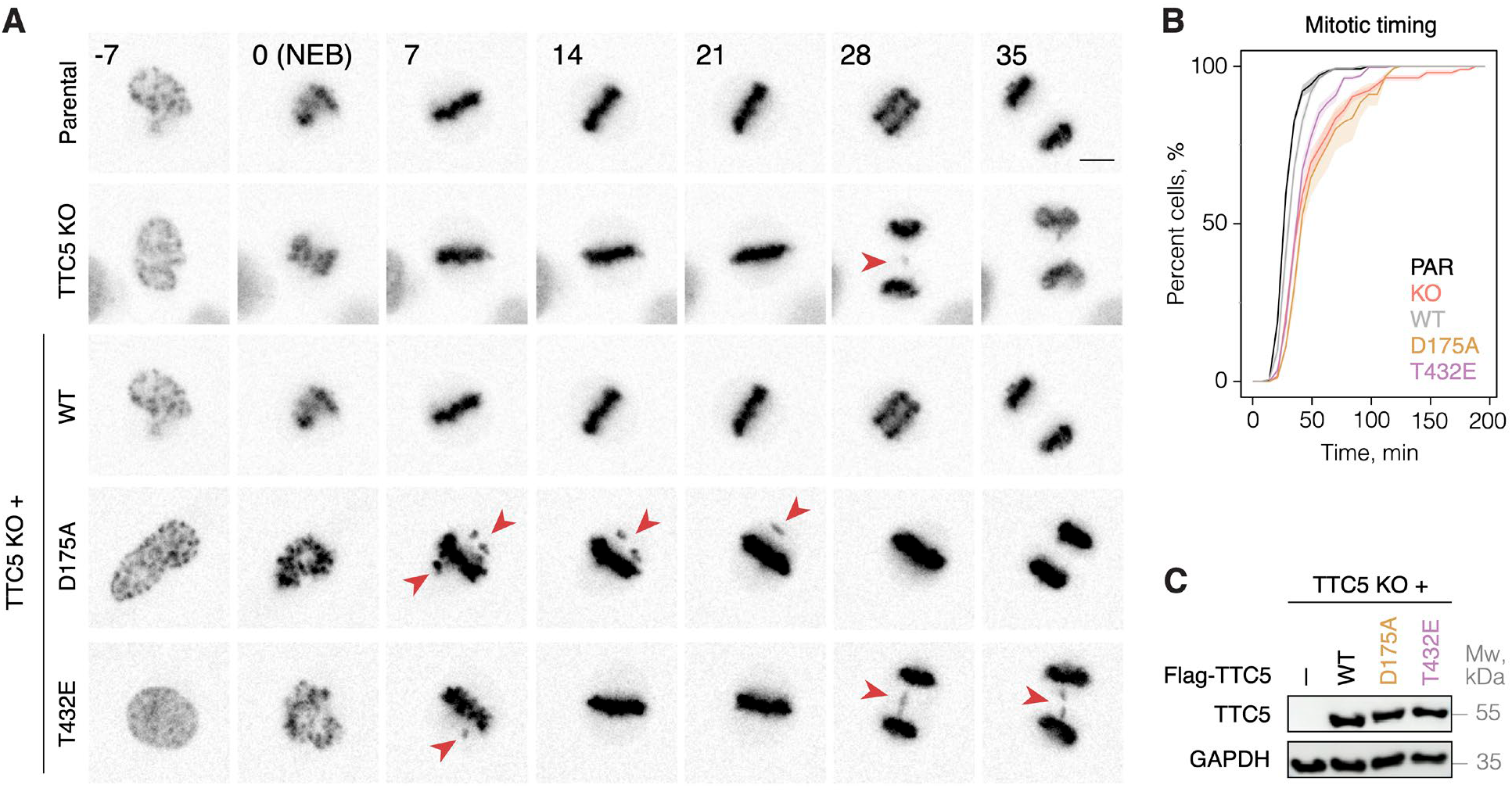
Loss of TTC5 repression by αβ-tubulins impairs mitotic fidelity (related to **Figure 4**). **(A)** Examples from live-cell imaging of HeLa parental, TTC5 knockout, and the indicated Flag-TTC5 rescue cell lines going through mitosis. DNA was visualized using SiR-DNA dye and maximum intensity projections of 3D volumes are shown. Frames were aligned to nuclear envelope breakdown (NEB, t = 0). Misaligned chromosomes in metaphase and segregation errors in anaphase are highlighted with red arrow heads. Scale bar = 10 μm. **(B)** Mitotic timing (time spent from NEB to anaphase onset) in HeLa cells of the indicated genotypes. Data are presented as cumulative frequencies ± SEM, from at least three independent replicates and more than 100 analyzed cells per genotype. **(C)** Total protein expression levels analysis by western blot for Flag-TTC5 constructs across the indicated cell lines used in panel A and in **Figure 4**.

